# Chronic Stability of Single-Channel Neurophysiological Correlates of Gross and Fine Reaching Movements in the Rat

**DOI:** 10.1101/674101

**Authors:** David T. Bundy, David J Guggenmos, Maxwell D Murphy, Randolph J. Nudo

**Author notes:** Corresponding Author: (RJN).

## Abstract

Following injury to motor cortex, reorganization occurs throughout spared brain regions and is thought to underlie motor recovery. Unfortunately, the standard neurophysiological and neuroanatomical measures of post-lesion plasticity are only indirectly related to observed changes in motor execution. While substantial task-related neural activity has been observed during motor tasks in rodent primary motor cortex and premotor cortex, the long-term stability of these responses in healthy rats is uncertain, limiting the interpretability of longitudinal changes in the specific patterns of neural activity during motor recovery following injury. This study examined the stability of task-related neural activity associated with execution of reaching movements in healthy rodents. Rats were trained to perform a novel reaching task combining a ‘gross’ lever press and a ‘fine’ pellet retrieval. In each animal, two chronic microelectrode arrays were implanted in motor cortex spanning the caudal forelimb area (rodent primary motor cortex) and the rostral forelimb area (rodent premotor cortex). We recorded multiunit spiking and local field potential activity from 10 days to 7-10 weeks post-implantation to characterize the patterns of neural activity observed during each task component and analyzed the consistency of channel-specific task-related neural activity. Task-related changes in neural activity were observed on the majority of channels. While the task-related changes in multi-unit spiking and local field potential spectral power were consistent over several weeks, spectral power changes were more stable, despite the trade-off of decreased spatial and temporal resolution. These results show that rodent primary and premotor cortex are both involved in reaching movements with stable patterns of task-related activity across time, establishing the relevance of the rodent for future studies designed to examine changes in task-related neural activity during recovery from focal cortical lesions.

## 1. Introduction

An important challenge in neuroscience is determining how the brain controls skilled forelimb movements, a topic that has important implications for motor recovery following brain injuries as well as the development of neuroprosthetic systems. Along with non-human primates, rodents are valuable models for examining the neurophysiological basis of motor control. In particular, rodents can learn to perform a wide variety of motor tasks, including: lever press/pull movements with complex timing [1], 2D center-out joystick movements [2], single-pellet reach-to-grasp food retrievals [3, 4], and even control of brain-computer interface systems with neural activity recorded from their motor cortex [5, 6]. Because of their ability to learn complex and flexible motor behaviors, rodent species have become valuable models for studies of motor control, neural plasticity during motor learning, and neural plasticity during recovery from a focal cortical injury [1, 7–11].

In humans and non-human primates, complex volitional movement is a result of the output and coordination of activity in several motor areas within the cortex. While not as extensive as the primate motor system, the wide range of motor behaviors examined in rodents is likely facilitated by the presence of multiple differentiated motor areas within the rodent motor cortex. The rodent motor cortex includes two distinct and interconnected areas in which forelimb movements can be elicited with intracortical microstimulation (ICMS) and which both have direct projections to the spinal cord: the caudal forelimb area (CFA) and the rostral forelimb area (RFA) [12, 13]. Based upon the sensory response properties and overall distribution of afferent and efferent fiber connections, it is thought that CFA is homologous to the primate primary motor cortex while RFA shows similarities to the premotor cortices and supplementary motor areas (SMA) [13, 14]. While CFA and RFA are functionally distinct, both demonstrate substantial task-related neural activity during the performance of a reaching task [7]. In part, this may be due to the dense reciprocal connectivity between the two regions [13], which depending on the relative timing of excitation in each area, allows RFA and CFA to modify the output from the other area [15].

In addition to contributing to the ability of rodents to perform complex motor behaviors, the presence of multiple distinct forelimb motor areas has important implications for studies examining neural plasticity in secondary motor regions during recovery from a cortical injury, such as stroke or traumatic brain injury. Following a lesion to CFA, rehabilitative training expands motor maps in RFA, suggesting that RFA plays some role in motor recovery [10]. However, the specific changes in the roles of CFA and RFA in controlling motor movements after a cortical injury, and the relevance of these changes to motor recovery, remain unclear. While examining task-related neural activity at different stages of motor recovery may help explain the observed anatomical and motor map changes during behavioral recovery, there are several considerations that need to be addressed prior to examining the specific correlates between changes in task-related patterns of neural activity and motor recovery. First, while single-pellet retrieval tasks involving reach-to-grasp movements are sensitive in measuring motor recovery [9, 10, 16, 17], more substantial lesion models can induce significant and persistent deficits limiting successful task performance for extended periods of time [9]. Therefore, a behavioral task with graded levels of difficulty will be required to assess the neural correlates of motor recovery across the full time course of motor recovery. Secondly, while previous studies have shown that chronic neurophysiological recordings can be acquired and used for decoding motor parameters and controlling brain-computer interface systems [5, 6, 11], the stability of the relationship between neural activity and motor movements at the level of individual channels is uncertain in rodent models.

This study addresses these limitations through a novel automated complex reaching task combining a ‘gross’ lever press with a ‘fine’ single pellet retrieval within a single trial. By combining these two components into a single task, it is possible to examine motor activity in a single animal while modulating the task requirements in terms of the level of fine motor control of the distal forepaw required to successfully complete the task. Additionally, we examined the stability of task-related neural activity over 7-10 weeks. Each task component was associated with robust task-related neural activity showing that both RFA and CFA are involved in controlling reaching movements. Furthermore, both multi-unit activity and local-field potentials were stable over periods of several weeks showing that these features can be used in future studies examining the longitudinal changes in movement-related neural activity during recovery from a cortical injury.

## 2. Materials and Methods

All procedures were approved by the University of Kansas Medical Center Institutional Animal Care and Use Committee in compliance with *The Guide for the Care and Use of Laboratory Animals* (Eighth Edition, The National Academies Press, 2011). To examine the neural correlates of gross and fine reaching movements, five Long-Evans rats (*Rattus norvegicus*) were trained to perform a novel complex reaching task utilizing a custom-designed automated behavior box while neural recordings were made from RFA and CFA.

### 2.1 Custom Behavior Box

A custom automated behavior box was developed that combined a lever press and skilled pellet retrieval into a single trial (Figure 1). The behavior box was constructed from acrylic sheets (12” × 12” × 18” tall, ¼” thickness). Each box had 15 mm wide vertical slits cut at 30 mm from the edge of the sides of both the front and back panels. A lever with an operating force of 20 g was placed directly behind each slit in the back panel with the paddle of the lever centered in the middle of the slit at a height of 28 mm, 23 mm behind the inside edge of the box. The position of the lever could be adjusted, allowing the lever to pass through the slit for temporary placement inside of the box to aid in the initial shaping of behavior. A shelf was placed spanning the entire width of the front of the box at a height of 30 mm. In front of the vertical slit, this shelf began 16 mm outside of the inner edge of the box and extended to 48 mm in front of the inner edge of the box, leaving a gap between the outer edge of the box and the front edge of the shelf. A depression (4 mm diameter, 1 mm depth) was made on the shelf 24 mm outside of the inside edge of the box and aligned to the inside edge of the vertical slit to ensure the consistent position of the food pellet in each trial. An acrylic door was placed in front of each slit in the front panel and was controlled by a linear actuator (Actuonix, Victoria, BC), controlling access to the shelf at the front of the box. Two infrared beam break sensors provided feedback regarding the position of the forepaw to aid in segmenting rodent behavior and switching between task periods. One infrared beam was placed across the front of the box and a second infrared beam was aligned vertically through the pellet location. Levers at the back corners of the box were coupled with the door at the diagonally opposite corner, forcing rats to use the same forepaw for each task component. By aligning the height and distance of the lever at the back of the box with the pellet tray at the front of the box, both the lever press and pellet retrieval required similar ballistic reaching movements with differing requirements for fine movements of the distal forepaw and sensorimotor integration. Task transitions were controlled by an Arduino microcontroller (Arduino Uno, Arduino, Ivrea, Italy) run by a custom-made MATLAB (Version R2017a, MathWorks, Natick, MA) executable function. A webcam was attached to the side wall, perpendicular to the front of the box, and aligned to image across the length of the shelf. Video from the webcam was acquired by the MATLAB executable code at 25 fps to capture reaches to the pellet tray and to control pellet dispensing. The webcam could be positioned at either side of the box to enable testing rats with either a right or left forepaw preference. Two pellet dispensers (Med Associates, Inc., Fairfax, VT) were used to dispense pellets to either side of the pellet shelf. Finally, an LED light was used to allow us to synchronize neural activity, behavioral performance, and videos recorded from additional external video camera.

**Figure 1.**
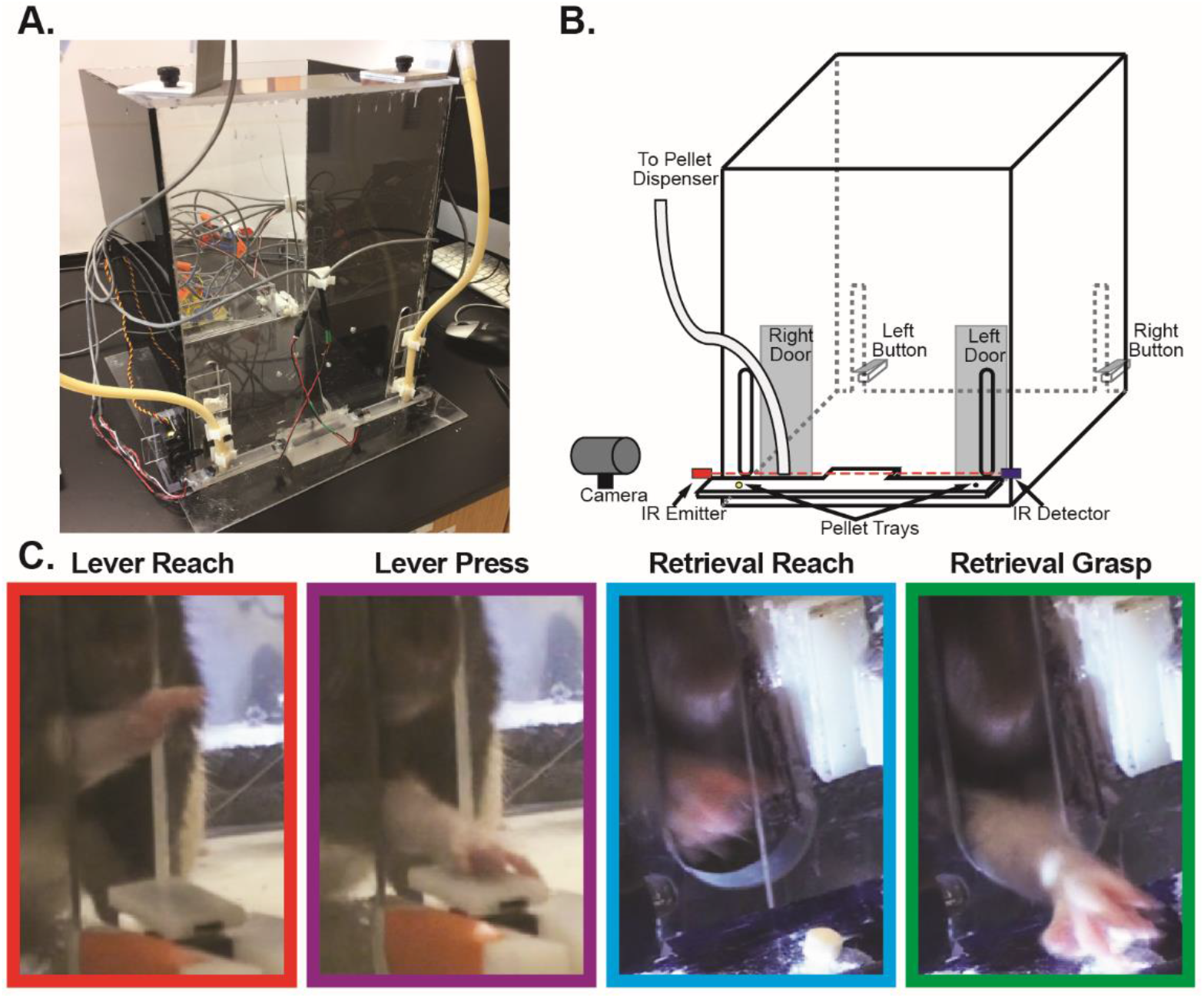
Complex Reaching Task. To evaluate the neurophysiological correlates of both gross and fine reaching movements within individual animals, we designed a novel, automated behavior box (**A**). **B.** At the beginning of each trial rats were required to use their preferred forearm to depress a lever placed outside an opening at the back corner of the box, causing a door at the front of the box in the corner diagonally opposite from the lever to open, providing access to a food pellet placed on a ledge outside the box. After transitioning to the door, rats reached through an opening to grasp and retrieve the food pellet reward. After detecting an attempt to retrieve the pellet via an infrared beam across the front of the box, the door was automatically closed, preventing repeated retrieval attempts. **C.** To examine the neurophysiological correlates of task performance, video recordings were used to identify the time points for the reach-to-button onset, button press, reach-to-pellet onset, and pellet grasp onset within each trial.

### 2.2 Task Structure

Prior to the beginning of each trial, both doors were closed, and a pellet was dispensed to the side of the pellet shelf corresponding to the rat’s preferred forelimb for pellet retrieval. Initially, rats were required to reach through the back of the box and depress the lever on the side of the box corresponding to their preferred forelimb. Upon depressing the lever, the door at the slit diagonally opposite the lever was opened, providing access to the previously dispensed food pellet on the shelf. After detecting a completed reach through the front of the box, as indicated by the infrared beam break being broken and then subsequently unbroken, the door was closed to limit secondary reaching attempts. If an attempt to retrieve the pellet was not made within 20 seconds, the trial was aborted.

### 2.3 Behavioral Training

Each rat went through a procedure to gradually shape their behavior to complete the full task. Initially, rats were placed in a non-automated box with a single slit cut in the center of the front panel. Rats were allowed free access to food pellets through this opening to learn to retrieve food pellets and determine forepaw preference. Pellets were placed in the center of the opening on a shelf placed across the front of the box at a height of 30 mm. Once rats began consistently retrieving pellets, their preferred forelimb was determined as the paw that was used on the majority of retrievals. Next, rats were introduced to the automated behavior box and allowed to retrieve pellets through the opening in the front of the box corresponding to their preferred forepaw. The door was closed after each retrieval attempt to familiarize rats with the door mechanism. The shelf in the automated behavior box had a gap between the edge of the shelf and the outer edge of the box which reinforced the requirement that the rats fully grasp each pellet, and not retrieve pellets by simply dragging them into the box. After successfully retrieving at least 50% of pellets, the lever press was incorporated into the training. Initially the front edge of the lever paddle was positioned inside the box and rats were cued to interact with the lever via auditory cues and food placed near the lever. A food pellet was dropped into the box by the secondary pellet dispenser each time the lever was depressed. This immediate reward served to reinforce the lever press. After beginning to depress the lever, the lever was gradually moved to its position for the full task, 23 mm outside the inside wall of the box. Next, we paired the lever press to the pellet retrieval. After each lever press, rats were cued to cross to the opposite corner of the box using auditory cues (tapping at the front corner of the box). After beginning to combine the lever press and pellet retrieval, the secondary reward received for the initial lever press was gradually extinguished to avoid any chewing artifact during the trial period. Finally, rats were trained until they successfully retrieved at least 50% of pellets in three consecutive training sessions with 25 trials per session. While total training time was variable, most rats completed each training step in 1-2 weeks and therefore could learn to adequately perform the entire task in 5-10 weeks with 30-minute long training sessions 3-5 days per week (mean ± SE = 33.6 ± 2.75 sessions). Following training, task performance was maintained with 2-3 training sessions per week until surgical procedures could be completed.

### 2.4 Surgical Procedures

Once rats achieved a threshold criterion of 50% successful retrievals on the full task in three consecutive sessions, we implanted chronic microwire electrode arrays into RFA and CFA. At the time of surgery, rats were 23-35 weeks old. Rats were initially anesthetized with isofluorane followed by injections of ketamine (80-100mg/kg) intraperitoneally and xylazine (5-10 mg/kg) intramuscularly. Rats were also given a preoperative dose of penicillin (45,000 IU subcutaneously) to limit the risk of infection. For the duration of the procedure, anesthetic state was confirmed by checking for the presence of a pinch reflex and corneal reflex, and a surgical level of anesthesia was maintained with supplemental injections of ketamine intramuscularly as necessary (10-20 mg/hr). Rats were placed in a stereotaxic frame and an incision was made along the midline of the scalp and the temporalis muscle was resected. A laminectomy was performed to reduce cortical swelling during the procedure. A craniectomy was then made over the sensorimotor cortex of the hemisphere contralateral to the preferred forelimb and the dura was retracted. Two 16-channel tungsten alloy micro-wire arrays with electrode diameters of 50 μm (Tucker-Davis Technologies, Alachua, FL) were implanted in each rat. Arrays were organized in a 2×8 grid with 250 μm spacing between electrodes and 500 μm spacing between rows. A silver wire from the probe was attached to a 00-80 stainless steel skull screw to act as a ground. The first electrode array was implanted into RFA with the second electrode array targeted to CFA as allowed by the size and orientation of the craniectomy and the rodent-specific pattern of vasculature within the craniectomy window. In three rats, the location of CFA and RFA were confirmed using ICMS mapping procedures as described previously [9]. In the remaining two rats, stereotaxic coordinates were used for RFA (3.5 mm anterior to bregma, 2.5 mm lateral to bregma) and CFA (0.5 mm anterior to bregma, 3.5 mm lateral to bregma) [9, 10, 18]. Each electrode array was implanted to a depth of approximately 1500 μm using a motorized micropositioner (Narishige International USA, Inc., Amityville, NY). An example ICMS map used to localize the implant locations is shown in Figure 2B. Electrode locations for all rats relative to skull landmarks are plotted in Figure 2C. The electrode locations in RFA overlap considerably across rats with some variability in the location of the CFA electrode arrays due to the limitations placed by the size of the craniectomy and vasculature pattern. After inserting the electrodes, the cortex was covered with a silicone elastomer (Kwik-Cast, World Precision Instruments, Sarasota, FL), a head cap was constructed from dental acrylic *in situ*, and the scalp was sutured around the head cap. Following the surgery, rats were given penicillin (45,000 IU subcutaneously) to limit the risk of infection. Four doses of buprenorphine (0.05-0.1 mg/kg subcutaneously) and acetaminophen (80-100 mg/kg orally) were given over the next 48 hours as analgesics. All rats were allowed to recover for 10-20 days prior to beginning neurophysiological recordings.

**Figure 2.**
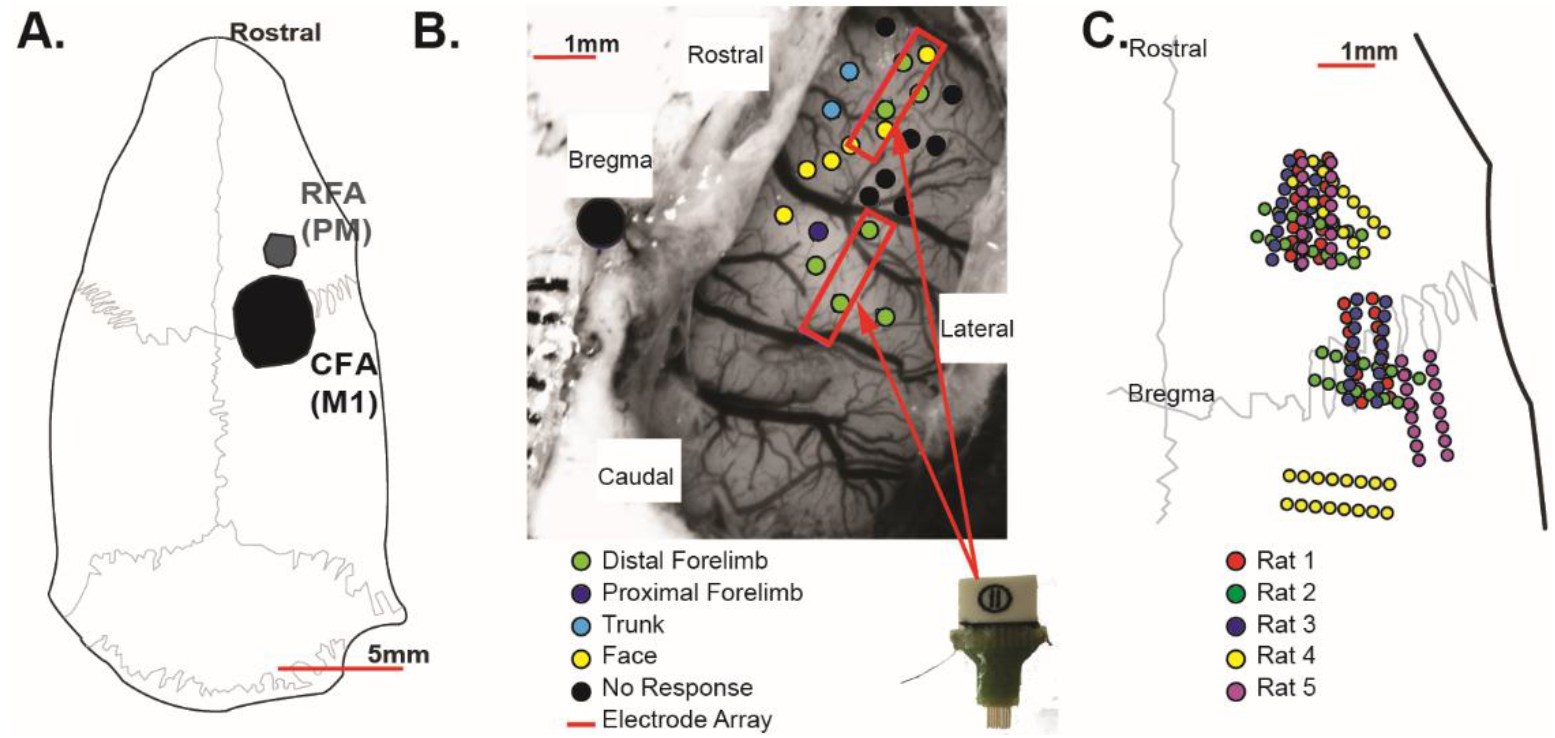
Chronic Microelectrode Implants. **A.** The rodent motor system consists of the caudal forelimb area (CFA), a homologue of M1, as well as a secondary rostral forelimb area (RFA), a homologue of premotor cortex. **B.** Sixteen-channel chronic microwire electrode arrays were implanted into each rat with one array placed in RFA and a second array implanted into CFA. Intracortical microstimulation mapping was used to confirm the locations of RFA and CFA in three rodents with steretotaxic coordinates used to determine implant locations in the remaining rats. **C.** Approximate implant locations relative to skull landmarks were determined from intraoperative photographs. With respect to stereotaxic coordinates, the location of the electrode arrays implanted into RFA was very consistent across rats with more variability in the locations of the CFA arrays due to the variable locations and orientations of blood vessels relative to the craniectomy opening.

### 2.5 Neurophysiological Recordings

Following the microelectrode implantation surgery, rats were given 10-20 days to recover from surgery. After the recovery period, we collected neurophysiological recordings while rats performed the reaching task in the automated behavior box. Rats performed the task 2-3 times per week for up to 10 weeks post-implant (mean ± SE = 14.6 ± 0.68 sessions). On recording days, rats were briefly anesthetized with isofluorane to ease connection of a recording headstage to each microelectrode array. The headstages, which performed amplification and digitization via an on-board amplifier chip (RHD 2132, Intan Technologies, Los Angeles, CA), were connected through a slip ring commutator (MC573, MOFLON TECHNOLOGY, Shenzhen, China) to an interface board (RHD2000, Intan Technologies, Los Angeles, CA) connected via USB to a PC computer. Neural activity was recorded at 20 kHz while rodents were in the automated behavior box performing the reaching tasks. For each session, signals were recorded while rats performed 45 trials or until 45 minutes had elapsed. Occasionally, recording sessions were truncated due to the headstages becoming unplugged prior to session completion.

### 2.6 Behavioral Scoring

During training sessions, time stamps of each lever press and infrared beam break were recorded by the recording amplifier as digital inputs. Videos of the reach-to-pellet were captured by a webcam at 25 fps and were synchronized to the neural recordings by the MATLAB interface. Additional video recordings were also made using an external digital video camera (Sony HDR-SR11, Sony Corporation, Tokyo, Japan) at 30 fps. The external camera was either positioned in front of the box to capture a higher resolution view of the reach to the food pellet, behind the lever to capture the lever press, or above the box to capture the rat’s movement throughout the box. This orientation alternated on each recording day. External videos were synchronized by capturing an LED light in the field of view that was controlled by the MATLAB interface and simultaneously recorded by the recording amplifier. Behavioral time points were co-registered by visually scoring the video recordings. As shown in Figure 1C, times for the reach to the lever, the lever press, the reach to the food pellet, and the grasp of the food pellet were identified. The times for the reach to the lever and reach to the food pellet were both defined as the frame in which the first forward movement of the paw to begin a reach was observed; the lever press was identified as the time that the lever circuit was electrically closed; and the grasp was defined as the first video frame where flexion of the digits at the end of the reach was observed. Successful trials were defined as trials in which the rat successfully retrieved the pellet on the first reach attempt to avoid potential confounding of neural activity related to an unsuccessful initial retrieval attempt followed by a successful second attempt. The reach to the food pellet and grasp were scored in each session using the webcam videos and confirmed using the external camera videos when available. While lever press time stamps could be identified in each recording, the reach to the lever was only identifiable in the subset of sessions with the external camera placed at the back of the box.

### 2.7 Data Analysis

Initially, raw local field potentials from all recordings were bandpass filtered from 2Hz-200Hz and visually screened to identify noisy or broken channels to exclude from all further analyses. Data was then processed to examine both multi-unit firing activity and local field potential (LFP) spectral power changes as described below.

#### 2.7.1 Multi-Unit Firing

To determine multi-unit spiking activity, signals were re-referenced to the common average of the non-noisy channels separately for each microelectrode array. Next, signals were band pass filtered from 300-3000Hz using an 8^th^ order elliptic filter. Potential spikes were identified based upon the more stringent criteria of either a fixed threshold of −50 uV or a variable monopolar threshold based upon the background amplitude of the signal for each channel [19]. Detected spikes were automatically clustered using an automated superparamagnetic clustering algorithm [19]. Manual spike sorting was then performed to eliminate non-physiological clusters. Figure 3A shows an example of bandpass filtered data from an exemplar recording session with a number of spike profiles. Figure 3C shows several spike profiles from an exemplar channel using the same recording as the traces shown in Figure 3A. Because we expected that it would be difficult to track single unit profiles across days, spike times from each cluster on a given channel were combined to yield the multi-unit activity for a given channel. Multi-unit firing rates were estimated by convolving a Gaussian waveform of one second duration and a standard deviation of 100 ms with the multi-unit spike times. To examine changes relative to baseline, 1000 random one-second long time windows, excluding all trial windows, were identified. The firing rate during these random time periods was estimated by convolving the multi-unit spike times with a Gaussian waveform as described above and the mean random firing rate was calculated by taking the mean across all time points from each random window. Finally, the firing rate during the recording was normalized by dividing the estimated firing rate by the mean random firing rate and log-transforming. The log-transform was used to normalize the firing rate with increases relative to baseline indicated by positive values and decreases relative to baseline indicated by negative values.

**Figure 3.**
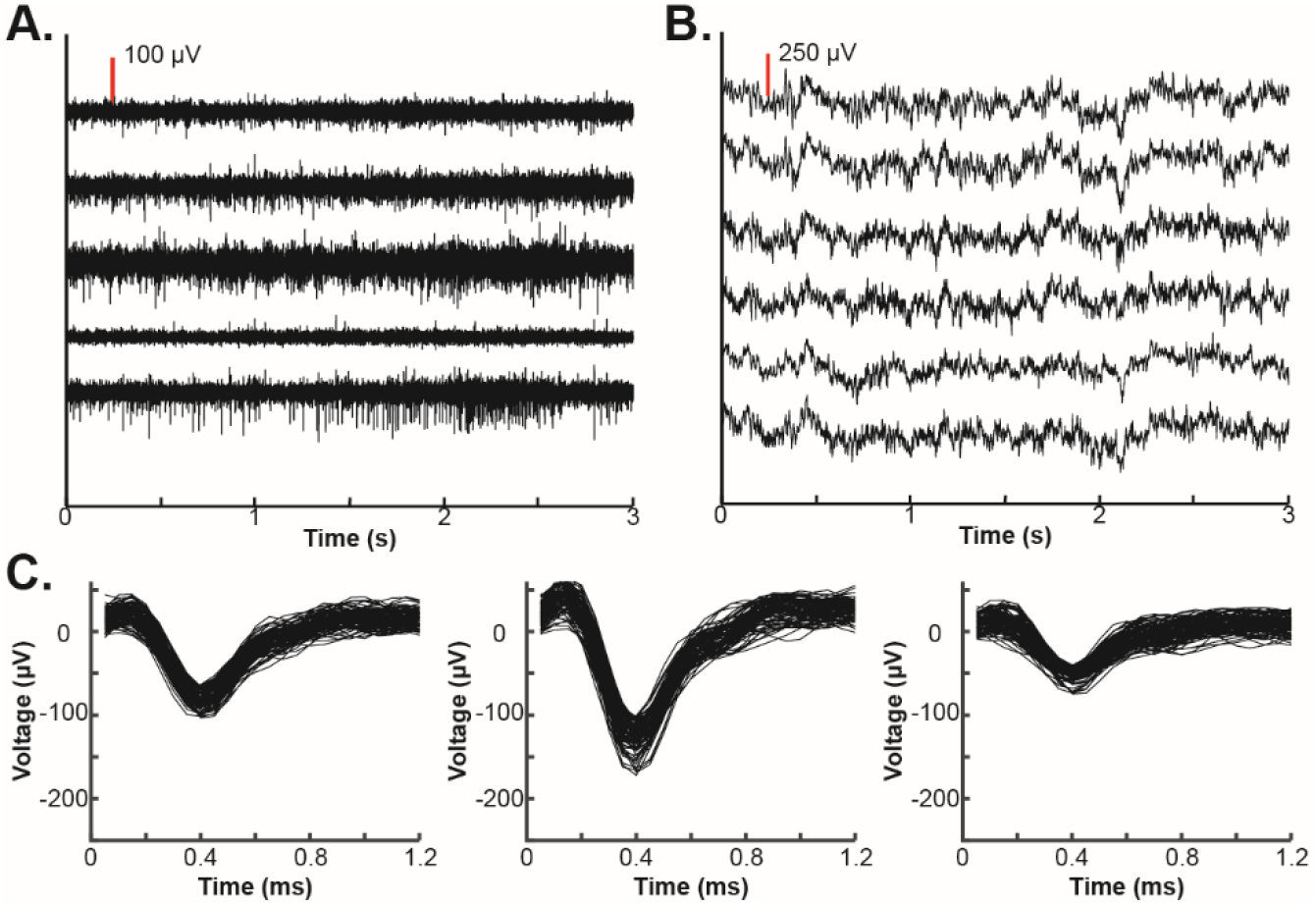
Electrophysiological Analyses. Multi-unit firing and local-field potential (LFP) activity were examined in each recording session. **A.** To examine multi-unit firing, signals were re-referenced to the common average and band-pass filtered between 300Hz and 3000Hz. Each trace shows data filtered for multi-unit activity and aligned to the same time point from randomly selected channels in a single rat. A scale of 100 μV is indicated by the vertical red line. **B.** LFP activity was visualized by filtering between 5Hz and 300Hz after re-referencing to the common average. Each trace shows data filtered for LFP activity and aligned to the same time point from randomly selected channels in a single rat. A scale of 250 μV is indicated by the vertical red line. **C.** To examine multi-unit firing, action potentials were detected using a superparamagnetic clustering algorithm followed by manual sorting to eliminate noisy clusters. Plots show 100 example spikes detected from 3 profiles isolated on a single channel. Following spike sorting, multi-unit activity was generated by combining all single-unit clusters from each individual channel.

#### 2.7.2 LFP Spectral Power Estimation

Task-related changes in the LFP spectral power were also examined. Raw signals were low-pass filtered at 400 Hz and decimated to a sampling rate of 1000Hz. All harmonics of the 60Hz power line noise below the Nyquist frequency were removed using an 8^th^-order Chebyshev notch filter. Signals were then re-referenced to the common average of all channels with neurophysiological signals separately for each microelectrode array. Examples of the raw LFP signals are shown in Figure 3B. The maximum entropy method, an autoregressive method of spectral estimation, was used to estimate spectral power [20]. A model order of 50 was selected and spectral power was estimated in 2 Hz frequency bins with bin centers ranging from 2 Hz to 200 Hz. Spectral power was calculated in 250 ms windows with shifts of 50 ms between windows to examine temporal changes. As with the multi-unit firing rate, 1000 random one-second long time windows, excluding all trial periods, were identified, and the spectral power was estimated in each window. Spectral power estimates were normalized by log-transformation and were then z-scored by subtracting the mean and dividing by the standard deviation of the frequency-specific spectral power estimated from the random time windows. Positive values indicated increases in spectral power at a given frequency relative to its baseline, and negative values indicated decreases in spectral power relative to baseline. Because the high-gamma band (70-105Hz) of the LFP signal has been found to represent localized activity that is strongly correlated with asynchronous spiking [21], the average high-gamma band power was found by averaging the z-scored spectral power for all frequency bins with centers between 70Hz and 105Hz.

### 2.8 Single-Day Characterization of Task-Related Neural Activity

Initially, the patterns of task-related neural activity during the two components of the task (lever press and pellet retrieval) were characterized using a single exemplar session. For the initial characterization of task-related neural activity, recordings were excluded for each rat until at least one session in which the rat performed 36 trials (80% of the daily goal) was acquired. To allow for examination of both components of the task, the next recording session with the external camera at the back of the box in which each rat performed at least 36 trials (80% of the daily goal) was used for the single-day characterization of task-related neural activity.

For each trial within the exemplar session, several behavioral time points were identified for further analysis. Sessions with the external camera behind the box allowed us to use the external camera recordings to identify the onset of reaching movements towards the lever and the lever timing to identify the downward movement used to press the lever, while the webcam at the front of the box was used to identify the onset of reaching movements to the food pellet and the onset of grasping movements. After identifying these time points for each trial, the task-related multi-unit firing rates and spectral power were examined as described below.

For each task event considered, a period from 1 s before to 1 s after the event was examined, matching the time of peak activations observed in previous studies of rodent reaching tasks [7]. For the lever press, task-related neural activity recorded from each channel was aligned either to the reach onset or lever press based upon whether the greatest absolute average depth-of-modulation was found by aligning neural activity to the onset of the reach towards the lever or to the lever press itself. Because the neural activity was normalized and log-transformed, signals were normally distributed with increases in neural activity relative to baseline indicated by positive values and decreases in neural activity indicated by negative values. Therefore, the depth-of-modulation was defined as the absolute value of the task-aligned neural activity. Similarly, for the pellet retrieval, each channel was aligned to the onset of the reach or onset of the grasp based upon whether the greatest absolute depth-of modulation was found by aligning neural activity to either the onset of the reach towards the pellet or the onset of the grasping movement. For the lever press, each channel was then classified as statistically significantly modulated and reach-related, statistically significantly modulated and press-related, or not significantly modulated. Similarly, for the pellet retrieval, each channel was also classified as significantly modulated and reach-related, significantly modulated and grasp-related, or not modulated. The statistical significance of each channel and task component’s classification was calculated to determine if the peak task-related neural activity was significantly different from chance using an independent samples t-test comparing the distribution of task-related neural activity at the time of the peak absolute depth-of-modulation to a randomly selected distribution of neural activity. The random distribution was derived by collecting the neural activity from 1000 random time points collected from outside of any trial period. A significance level of p<0.05 was used to define significance with Bonferoni correction for the total number of comparisons tested across channels and task periods. Each channel with spike profiles identified was classified as described above. Each channel was separately classified based upon whether there were statistically significant increases or decreases of the high-gamma band spectral power during either the lever press or the pellet retrieval. As with the multi-unit firing, the high-gamma band was also used to give each channel two classifications: first, each channel was classified as statistically significantly modulated and reach-related, statistically significantly modulated and press-related, or not significantly modulated for the lever press, and second, each channel was classified as statistically significantly modulated and reach-related, statistically significantly modulated and grasp-related, or not significantly modulated for the pellet retrieval.

### 2.9 Chronic Stability of Task-Related Neural Activity

Finally, we sought to examine the chronic stability of task-related neural activity in RFA and CFA. Because videos capturing the lever press were not captured for every recording day due to rotating the position of the external video camera, the analysis of the stability of task-related neural activity was limited to the pellet retrieval component of the task. For each channel with a statistically significant reach or grasp-related change in multi-unit firing rate in the exemplar recording session described above, trials from each recording day were aligned to the reach onset or grasp onset based upon the initial classification from the exemplar recording day. Next, the daily average multi-unit firing rate was calculated for each channel. A global average time course of task-aligned multi-unit firing rate was calculated by averaging the firing rate from all trials across all recording days. Finally, for each channel and recording day, the correlation coefficient (Pearson’s r) was calculated between the global and daily averaged firing rate. The consistency of task-related high-gamma band power changes was assessed using the same procedure for all channels with statistically significant reach or grasp-related changes in high-gamma band spectral power in the exemplar session. We then compared the inter-day consistency of single-channel task-related multi-unit firing rate changes with the inter-day consistency of task-related high gamma band power changes using a Wilcoxon rank sum test comparing the distribution of correlation values for the two signal types across all rats, channels, and days.

## 3. Results

### 3.1 Rat Characteristics and Behavioral Performance

All rats learned to perform the task with successful retrievals on 60-80% of trials. Table 1 summarizes the overall behavioral performance of each rat. Each rat continued to perform the task with accuracies above 50% success rates, indicating that the microelectrode implantation did not impair the ability to perform the task (Exemplar Sessions: mean ± SE = 67.4% ± 4.0%; Overall: mean ± SE = 63.5% ± 1.4%). While these performance scores are lower than reported in previous studies of pellet retrieval in rats [9, 10, 17], the gap between the shelf and box required rats to fully lift each pellet and the door restricted repeated reaching attempts, increasing the difficulty of the task. Additionally, to better isolate task-related neural activity, successful trials were defined as trials in which the rat successfully retrieved the pellet on the first attempt. The accuracies observed were similar to other studies with similar placements of pellets on a pedestal, increasing the chances of the rat dislodging the pellet off of the shelf, or in which reaching success was scored using the first attempt [3, 22, 23]. Microelectrode implant locations relative to skull landmarks are shown in Figure 2C. While microelectrodes were implanted contralateral to the preferred forepaw, microelectrode locations in the left hemisphere were reflected across the midline so all microelectrode locations could be visualized on the right hemisphere to allow for comparison across animals. Because we prioritized placing the first array into RFA given its smaller size, the locations of the RFA arrays were highly consistent across rats. The increased variability in the location of CFA arrays is due to the geometric limitations imposed by the craniectomy orientation, patterns of vasculature, and size of the microelectrode arrays as opposed to variability in the location of CFA across rats. While the position of the CFA microelectrode arrays varied, the majority of the individual electrode wires were still within the CFA motor representation.

**Table 1.** Summary of Task Performance.

| Rat | Preferred Forepaw | Total Sessions | Total Trials | Successful Trials | Percent Accuracy | Exemplar Session Date (Days Post-Implant) | Number of Exemplar Trials | Successful Exemplar Trials | Exemplar Percent Accuracy |
| --- | --- | --- | --- | --- | --- | --- | --- | --- | --- |
| 1 | Right | 13 | 371 | 215 | 58.0% | 25 | 36 | 23 | 63.9% |
| 2 | Right | 13 | 506 | 332 | 65.6% | 24 | 44 | 28 | 63.6% |
| 3 | Left | 16 | 671 | 440 | 65.6% | 13 | 44 | 32 | 72.7% |
| 4 | Left | 16 | 648 | 411 | 63.4% | 20 | 45 | 36 | 80.0% |
| 5 | Left | 15 | 493 | 320 | 64.9% | 24 | 44 | 25 | 56.8% |

### 3.2 Task-Related Neural Activity During ‘Gross’ and ‘Fine’ Reaching Movements

Initially, we sought to characterize the patterns of task-related neural activity observed in an exemplar recording session. The task-related multi-unit firing activity from several exemplar channels is shown in Figures 4-6. In the first channel (Figure 4A and C), a statistically significant increase in firing rate is observed during reaching in both task contexts. Specifically, the peak multi-unit firing was significantly different from chance when aligned either to the reach to the lever or the reach towards the pellet, with a stronger modulation observed during the pellet retrieval. Additionally, when compared to the pre-movement period, there was a weaker but consistent increase in firing rate maintained during the period between the lever press and the pellet retrieval. While this period was variable in length, the maintained increase in neural activity is apparent either when aligned to the reach towards the lever or the reach towards the pellet. In a second exemplar channel (Figure 5A and C) a grasp-related increase in firing rate is illustrated. In this channel, there was a small but statistically significant increase in firing rate around the lever press and a larger statistically significant increase in firing rate observed when aligned to the grasp. While this increase began before the reach towards the pellet, there was a stronger depth-of-modulation observed when aligning trials to the grasp than when aligning trials to the reach onset. Finally, other channels showed a more complex response. For example, the channel shown in Figure 6 displayed a statistically significant decrease in firing rate with a peak just after the lever press and a statistically significant increase in firing rate immediately before the reach towards the food pellet.

**Figure 4.**
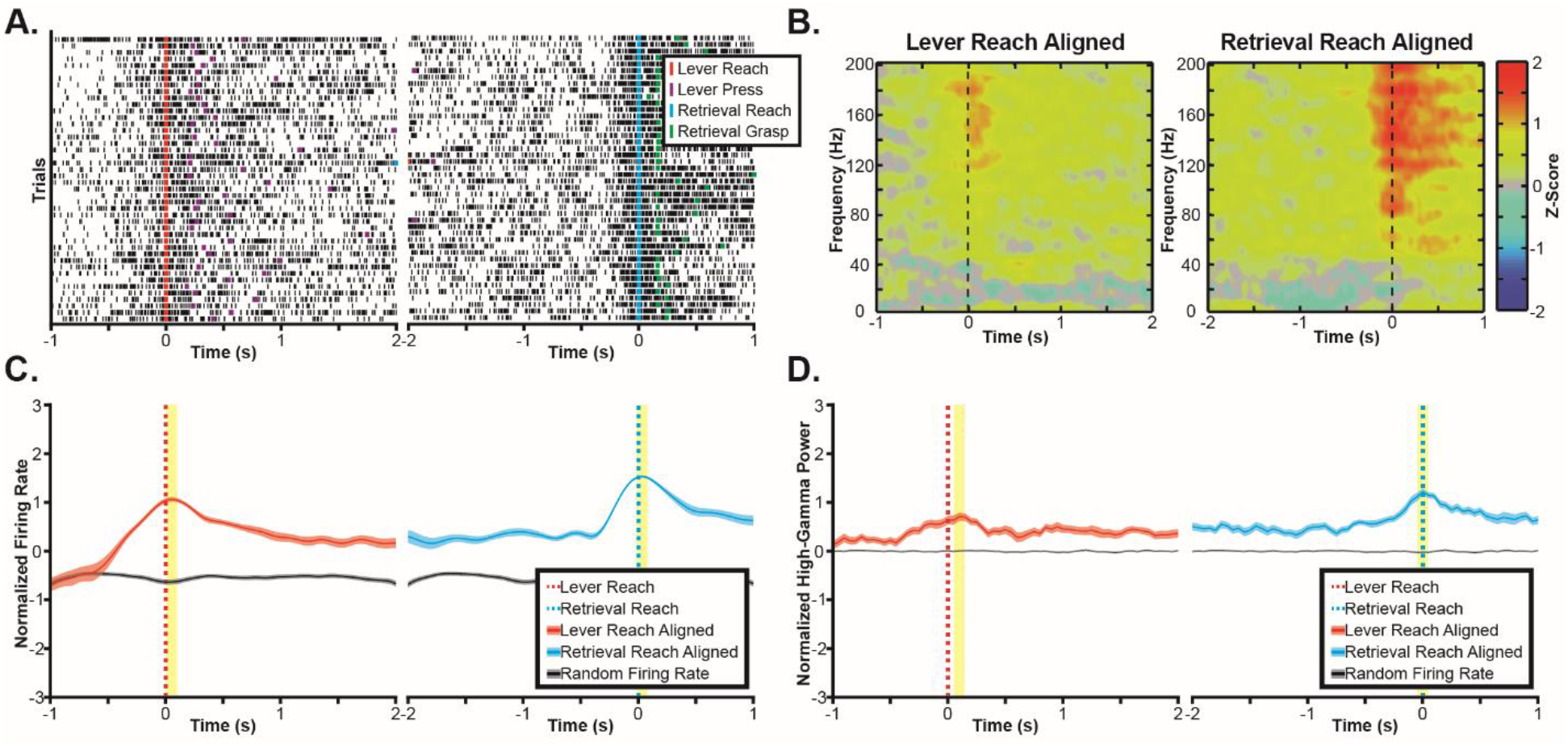
Exemplar Reach-Related Channel. **A.** Raster plots show multi-unit spike times (black ticks) aligned to the onset of the reaching movement towards the lever (left) and food pellet (right), showing an increase in firing rate correlated to the reach onset. **B.** Spectral power changes are plotted aligned to the onset of reaching movements to the lever (left) and pellet (right). Spectral power changes were log transformed and z-scored relative to randomly selected time windows between trials, therefore, positive values indicate increases in spectral power, and negative values indicate decreases in spectral power. A broadband increase in spectral power in frequencies above 60Hz was observed for both the reach to the button and the reach to the pellet. **C.** Multi-unit firing rates were estimated by convolving a Gaussian waveform with spike times and then averaging firing rates across trials, showing a statistically significant (p<0.05) increase in firing rate aligned to the onset of the reach in both task components. The statistically significant peak firing rates are indicated by the yellow highlighting. **D.** High-gamma band (70Hz-110Hz) power also significantly increased around both the reach to the lever (left) and the reach to the food pellet (right). The statistically significant peak modulations of high-gamma band spectral power are indicated by the yellow highlighting.

**Figure 5.**
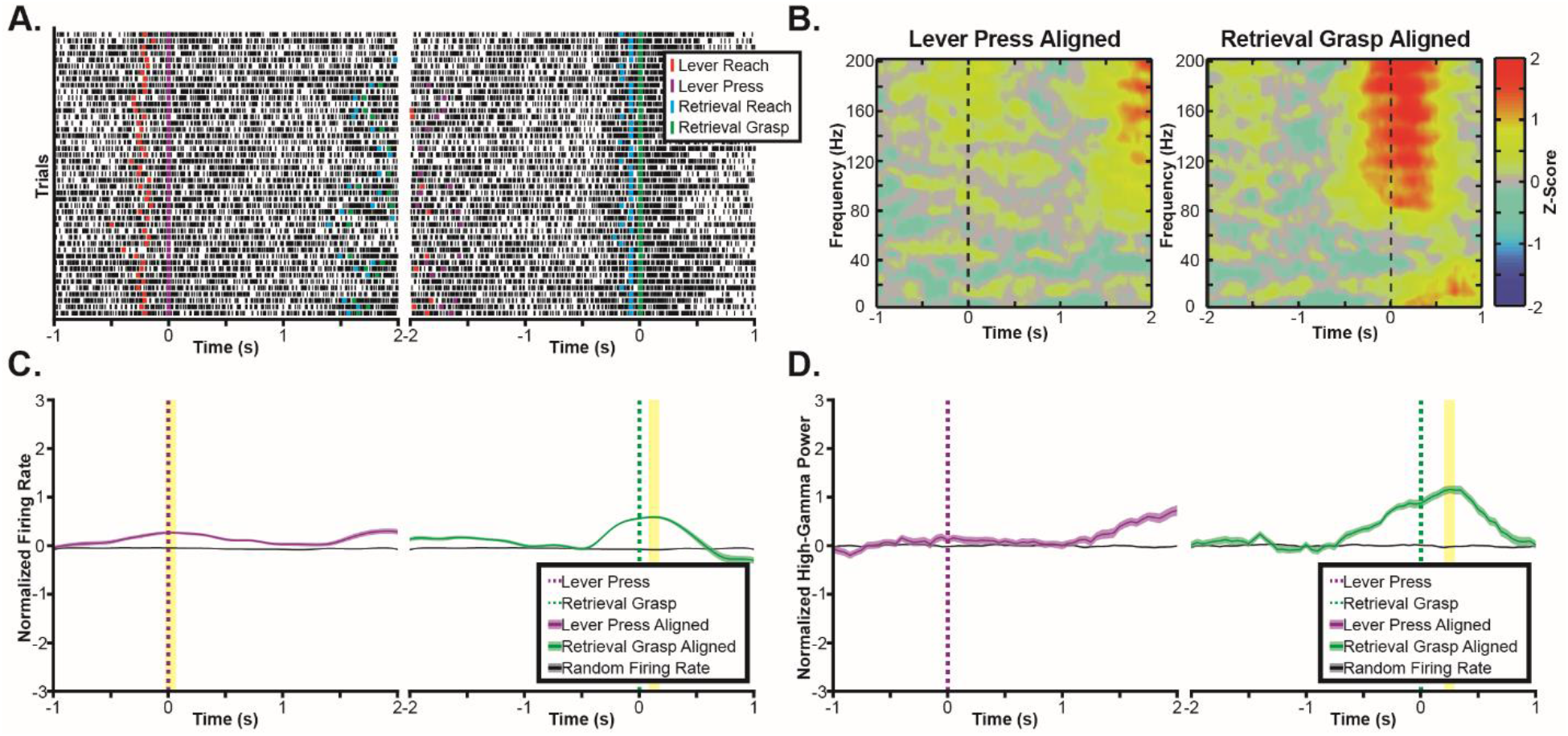
Exemplar Grasp-Related Channel. **A.** Raster plots show multi-unit spike times (black ticks) aligned to the lever press (left) and onset of the pellet grasp (right), showing an increase in firing rate correlated to the grasp onset. **B.** Spectral power changes are plotted aligned to the lever press (left) and pellet grasp (right). Spectral power changes were log transformed and z-scored relative to randomly selected time windows between trials, therefore, positive values indicated increases in spectral power, and negative values indicate decreases in spectral power. A broadband increase in spectral power in frequencies above 60Hz was observed for the pellet grasp. **C.** Multi-unit firing rates were estimated by convolving a Gaussian waveform with spike times and then averaging across trials. Mean firing rates synchronized to the grasp were significantly (p<0.05) higher than expected by chance. The statistically significant peak firing rates are indicated by the yellow highlighting. **D.** High-gamma band (70Hz-110Hz) power also significantly increased around the pellet grasp (right). The statistically significant peak modulation of high-gamma band spectral power during the grasp is indicated by the yellow highlighting.

**Figure 6.**
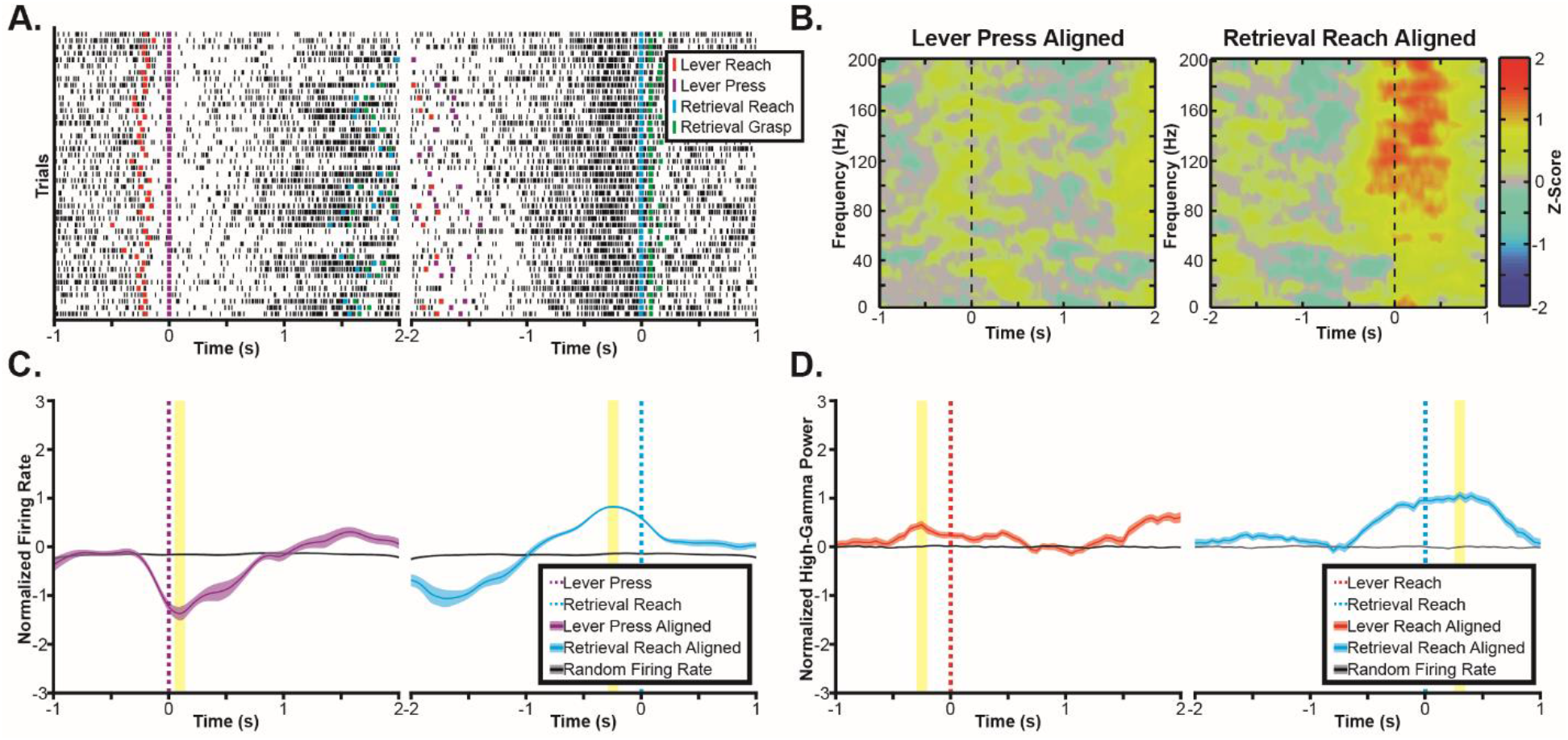
Exemplar Pellet-Related Channel. **A.** Raster plots show multi-unit spike times (black ticks) aligned to the lever press (left) and onset of the reach to the pellet (right), showing a decrease in firing rate correlated to the lever press followed by an increase in firing rate prior to the reach to the pellet. **B.** Spectral power changes are plotted aligned to the reach to the lever (left) and reach to the pellet (right). Spectral power changes were log transformed and z-scored relative to randomly selected time windows between trials, therefore, positive values indicated increases in spectral power, and negative values indicate decreases in spectral power. A broadband increase in spectral power in frequencies above 60Hz was observed for the reach to the food pellet (right) with a smaller increase in broadband spectral power observed before the reach to the lever. **C.** Multi-unit firing rates were estimated by convolving a Gaussian waveform with spike times and then averaging across trials. Mean firing rates show a complex task response with firing rates significantly (p<0.05) decreased around the lever press and then significantly (p<0.05) increased immediately prior to the reach to the food pellet. The statistically significant peak firing rates are indicated by the yellow highlighting. **D.** High-gamma band (70Hz-110Hz) power did not show a decrease around the lever press but instead showed a small, but statistically significant (p<0.05), increase in high gamma band power prior to the reach to the lever. The high-gamma band power maintained the significant increase in activity around the reach to the food pellet observed in the multi-unit firing rate with an increased duration of activity. The statistically significant peak modulations of high-gamma band spectral power are indicated by the yellow highlighting.

Along with modulations of multi-unit firing rate, widespread modulations of the high-gamma band spectral power were also observed. Figures 4-6 also contain time-frequency plots showing task-related changes in spectral power throughout the frequency spectrum as well as the specific change in high-gamma band (70-105Hz) power for the same exemplar channels as the multi-unit firing rate. Across each exemplar, increases in high-gamma band power were observed in the same channels and task periods where increases in multi-unit firing rate were observed. Specifically, increases in broadband spectral power were observed when aligned to both the reach to the lever and reach to the pellet in the channel shown in Figure 4, for the grasp but not the lever press in the exemplar channel shown in Figure 5, and for the reach to the pellet in Figure 6. While increases in multi-unit firing rate and high gamma band power were often observed in the same channel, the high gamma band did not exhibit task-related decreases. In the exemplar channel shown in Figure 6, firing rate decreased around the lever press, while a small but statistically significant increase in high-gamma band spectral power was observed. The time scales of changes in high-gamma band spectral power were also often extended relative to multi-unit firing rate. This longer modulation of activity is particularly apparent for the pellet retrieval in Figure 5 (grasp) and Figure 6 (reach to the pellet).

The range of task-related activations observed was characterized across all channels and all animals. The topography of microelectrodes classified as related to each task component is shown in Figure 7. The proportion of channels classified as related to each task component are summarized for multi-unit firing rate changes in Table 2. Of the 118 channels with at least one spike profile found, 98 channels had statistically significant task-related changes in multi-unit firing rate. While more channels had firing-rate changes during the pellet retrieval than the lever press, more channels were classified as reach-related than were classified as lever press-related or grasp-related. For the pellet retrieval this difference in the number of reach-related and grasp-related channels appears to stem from RFA, where almost three times more channels were classified as reach-related than were classified as grasp-related. The proportion of channels classified as related to each task component using high-gamma band power changes are shown in Table 3. Because LFP power changes can be detected in the absence of spike profiles, more channels had statistically significant changes in high gamma band power than had changes in multi-unit firing rate. For lever press high-gamma band power changes, the vast majority of channels were classified as reach-related, with a smaller number classified as lever press-related. Classification using high-gamma band spectral power was similar to the classification using multi-unit firing activity for the lever press. However, this classification differed for pellet retrieval. When classified based upon the changes in high-gamma band spectral power, the proportion of reach and grasp-related channels were similar, with a slightly higher number of reach-related than grasp-related channels in RFA.

**Figure 7.**
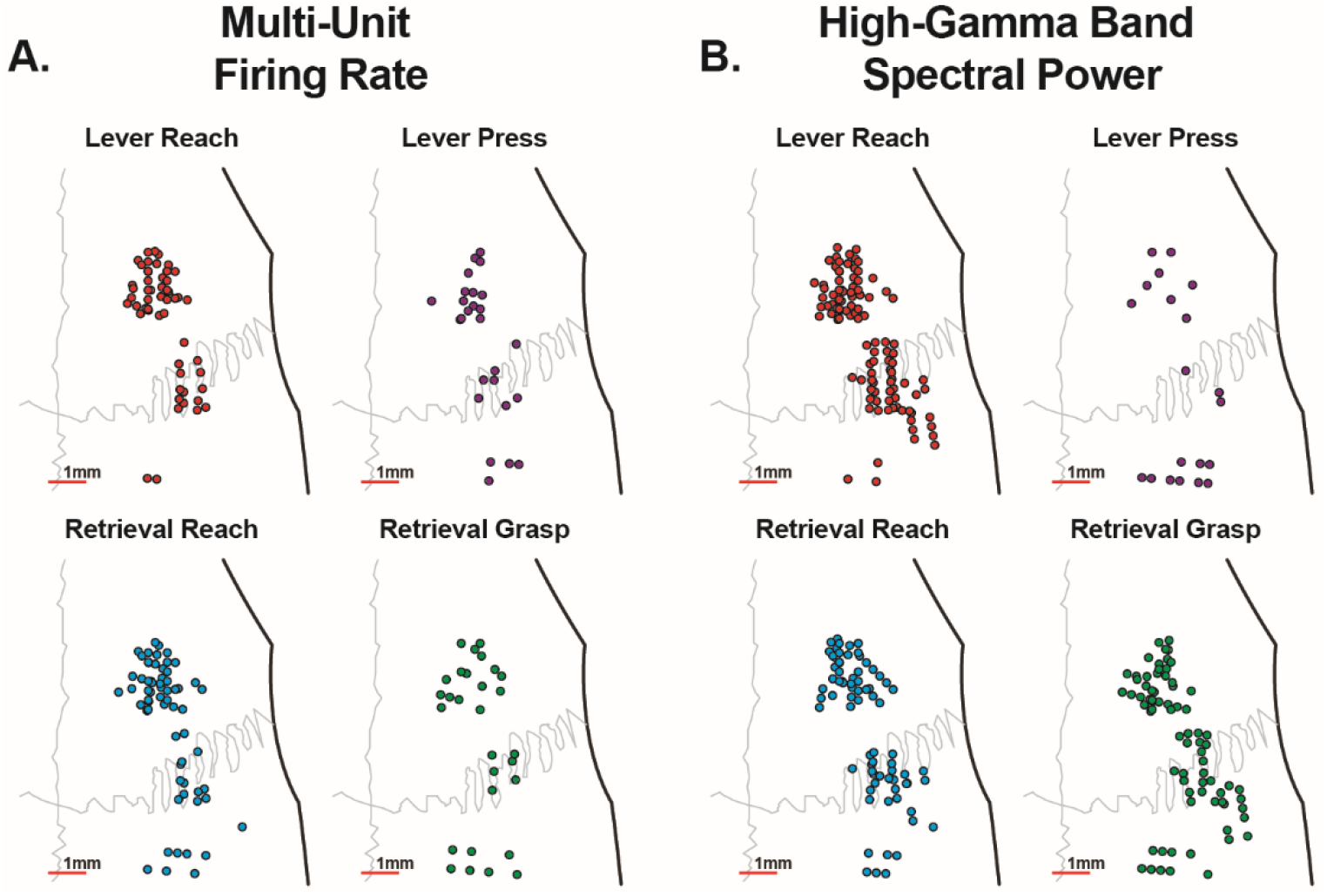
Microelectrode Classification. Each microelectrode was classified based upon whether it was significantly modulated by the reach to the lever, the lever press, or not modulated during the lever press. Similarly, each electrode was also classified as modulated when aligned to the reach to the food pellet, when aligned to the grasp of the pellet, or not modulated during the retrieval. Each classification was made using both the multi-unit firing rate (**A**) and high-gamma band (70-110Hz) spectral power (**B**). When examining changes in multi-unit firing rate, more channels were modulated by the reaching movements than either the lever press or grasping movements. Additionally, more of the channels modulated by reaching movements were located in RFA than in CFA. When examining changes in high-gamma band power, while more channels were modulated by the reach than the button press, because of the decrease in temporal specificity, a similar number of channels were modulated by both the reach and the grasp. In contrast to the multi-unit firing rate, similar numbers of electrodes with significant task-related activations aligned to the reach were found in both CFA and RFA.

**Table 2.**
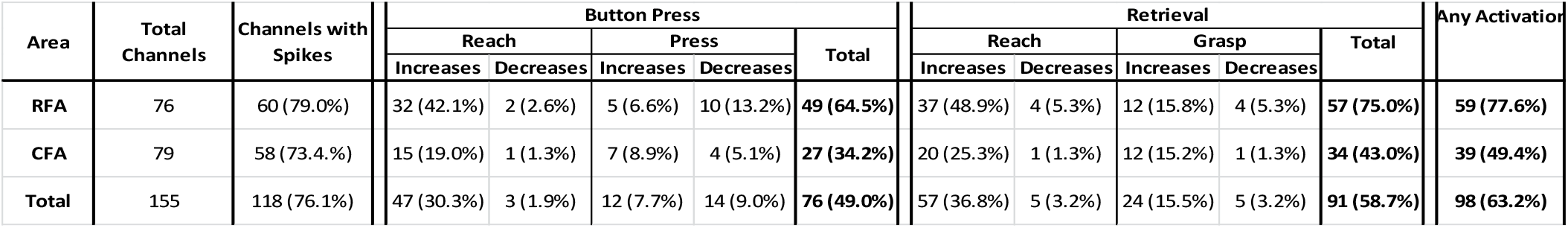
Summary of significant task-related changes in multi-unit firing rate.

**Table 3.** Summary of significant task-related changes in high-gamma band (70-110Hz) power.

| Area | Total Channels | Button Press |  |  | Retrieval |  |  | Any Task Modulation |
| --- | --- | --- | --- | --- | --- | --- | --- | --- |
|  |  | Reach | Press | Total | Reach | Grasp | Total |  |
| RFA | 78 | 53 (68.0%) | 8 (10.3%) | **61 (78.2%)** | 41 (52.6%) | 37 (47.4%) | **78 (100%)** | **78 (100%)** |
| CFA | 79 | 50 (63.3%) | 12 (15.2%) | **62 (78.5%)** | 33 (41.8%) | 39 (49.4%) | **72 (91.1%)** | **73 (92.4%)** |
| **Total** | **157** | **103 (65.6%)** | **20 (12.7%)** | **123 (78.3%)** | **74 (47.1%)** | **76 (48.4%)** | **150 (95.5%)** | **151 (96.2%)** |

### 3.3 Consistency of Rodent Task-Related Neural Activity

In addition to characterizing the task-related activations observed in RFA and CFA, we investigated whether task-related activations in RFA and CFA were stable over several weeks. The task-related multi-unit firing and spectral power changes for an exemplar channel are shown for several recording sessions in Figure 8. On each recording day, an increase in multi-unit firing rate was observed around the reach towards the pellet. Because the background firing rate differed across days, the depth-of-modulation was variable across days. In contrast, a similar broadband increase in high-gamma band power was observed for each session with a similar depth-of-modulation observed in each day shown.

**Figure 8.**
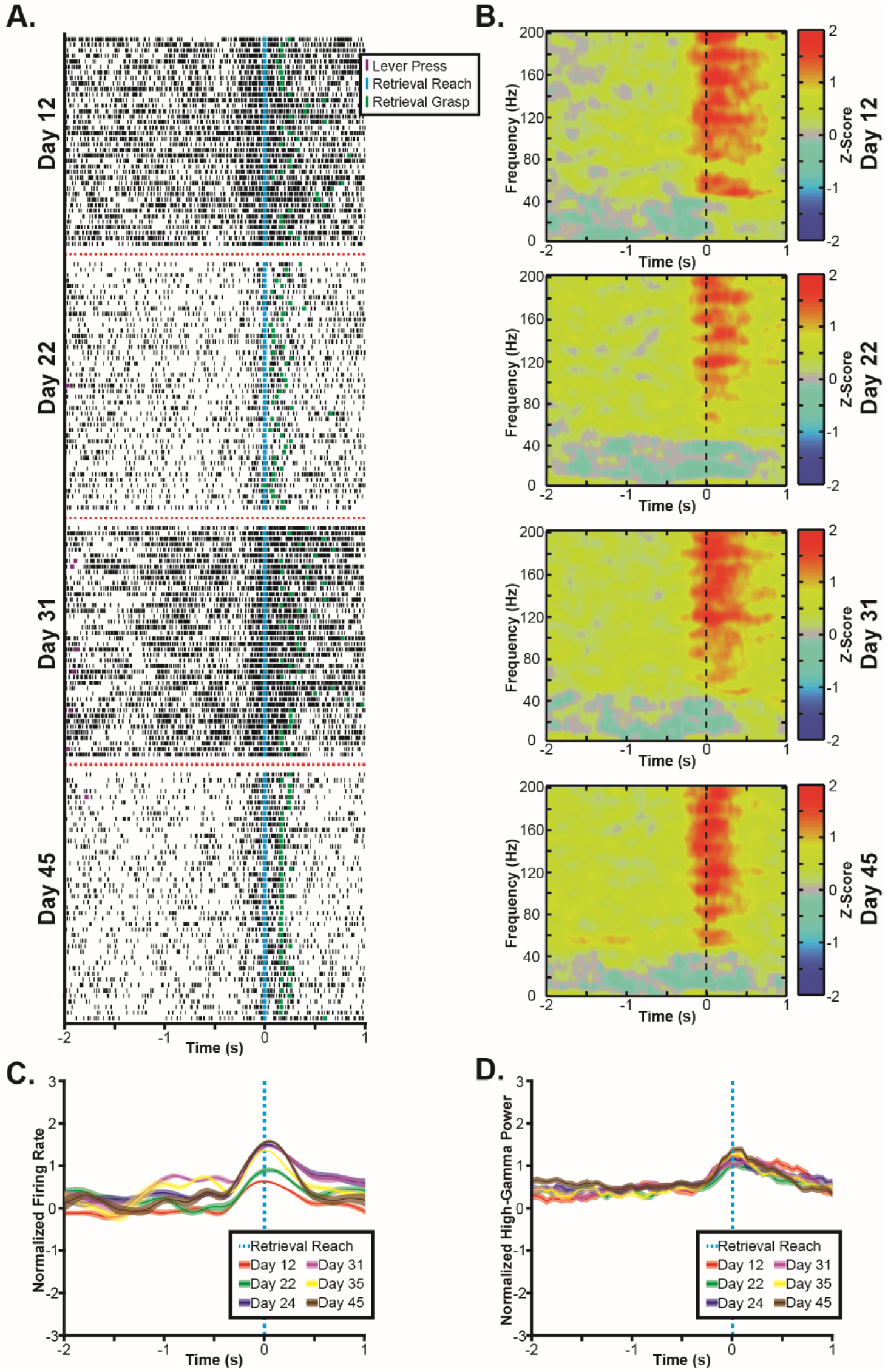
Exemplar task-related activity across recording days. **A.** Raster plots show multi-unit spike times (black ticks) aligned to the onset of the reaching movement towards the food pellet on several recording days, showing an increase in firing rate correlated to the reach onset that is conserved across days. **B.** Spectral power changes are plotted aligned to the onset of reaching movements to the pellet on the same days. Spectral power changes were log transformed and z-scored relative to randomly selected time windows between trials, therefore, positive values indicate increases in spectral power, and negative values indicate decreases in spectral power. A broadband increase in spectral power in frequencies above 60Hz is observed aligned to the reach in each day. **C.** Multi-unit firing rates were estimated by convolving a Gaussian waveform with spike times and then averaging firing rates across trials, showing a statistically significant (p<0.05) increase in firing rate aligned to the onset of the reach. While the timing of this increase in firing rate is conserved across days, there is some variance in the depth-of-modulation observed. **D.** High-gamma band (70Hz-110Hz) power also significantly increased around the reach to the food pellet with similar depth-of-modulations observed on each recording day. All plots were generated from the same exemplar channel within RFA of a single animal.

The stability of task-related neural activity was quantified by calculating the correlation between the daily and overall average modulation for each channel with a significant task-related modulation in the exemplar recording session. These correlations are shown for each rat in Figure 9 and summarized in Table 4. The task-related neural activity was largely stable for each rat, decreasing only when recording quality becomes impaired, such as in Rat 1 starting around day 70 post-implant and in Rat 2 starting around 40 days post-implant. While both multi-unit firing rate changes and high-gamma band power changes were stable over time, daily task-related changes in high-gamma band power were more correlated to the overall average task-related changes in high-gamma band power than multi-unit firing rates were. Specifically, the distribution of correlations between daily and overall task-related modulations was significantly higher for high-gamma band spectral power changes than for multi-unit firing rate changes (Wilcoxon rank sum test, p=4.04 × 10^−44^).

**Figure 9.**
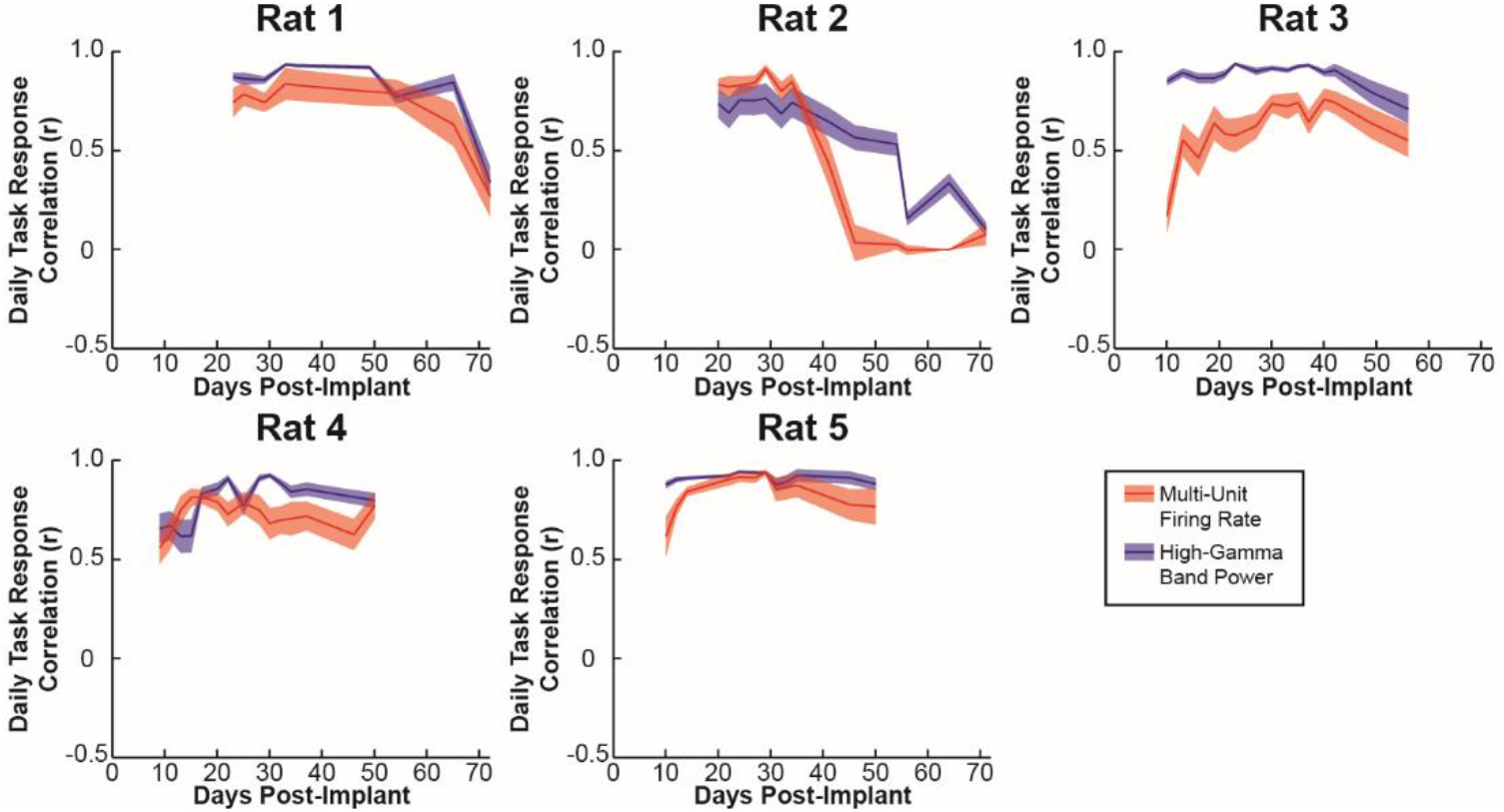
Cross-Day Stability. The stability of task-related changes in neural activity was examined by determining the correlation between the daily mean firing rate and the overall mean firing rate as well as the correlation between the daily high-gamma band power change and the overall mean high-gamma band power change for each channel. Plots show the mean daily correlation across channels and the error bars show the standard error. While a decrease in signal quality was observed at 9 weeks and 5 weeks in rats 1 and 2 respectively, overall, task-related changes in multi-unit activity and high-gamma band power were stable across days for each rat. While changes in multi-unit firing rate and high-gamma band power were both stable across recording days, the task-related the high-gamma band power change had higher correlations between the daily and overall mean than was observed for task-related multi-unit firing rate changes. Additionally, high-gamma band power changes show a slower decrease in stability from the degradation in signal quality observed in rats 1 and 2.

**Table 4.**
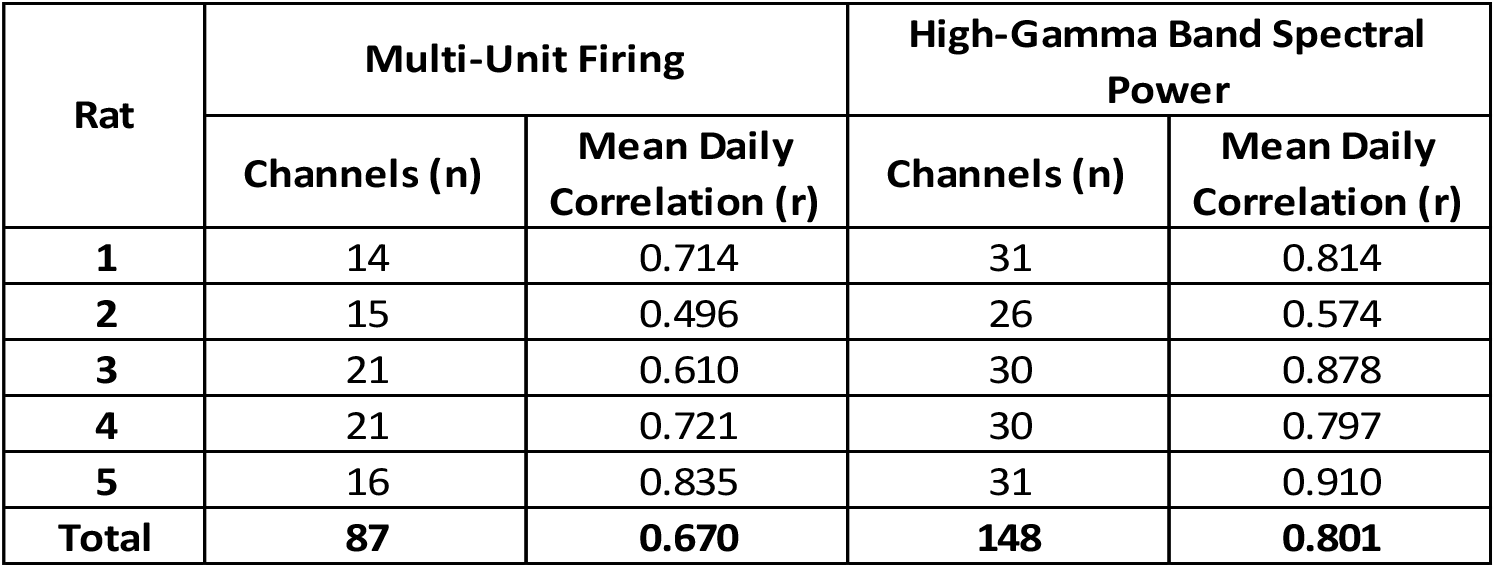
Summary of the stability of task-related changes in multi-unit firing rate and high-gamma band power.

## 4. Discussion

This study used a novel automated behavior box to demonstrate the widespread nature of reach and grasp-related neural activity in the rat primary motor cortex (CFA) and premotor cortex (RFA). Previous studies have demonstrated task-related modulation of activity during reaching tasks including center-out movements of a lever, push/pull movements of a lever, and pellet retrieval tasks [1, 7, 11, 24–26]. Importantly, these previous studies found that neural activity could be used to decode movement parameters, such as kinematics and kinetics [11, 25]. Here we examined the neural correlates of reaching movements using a novel automated behavior box that combined a gross reaching task (lever press) and a fine reaching task (skilled pellet retrieval) in close temporal succession. While previous examples of automated rodent behavior boxes have been described, our system adds the ability to examine two similar but distinct and relatively unconstrained reaching tasks in the same animals [22, 23, 27]. As with these previous systems, the use of an automated system reduces the level of supervision necessary to train rodents to perform reaching movements and eliminates many potential biases inherent in manually administered tasks.

Interestingly, a greater number of channels were active during the skilled pellet retrieval than during the lever press, potentially indicating greater cortical involvement in planning and executing the pellet retrieval than the lever press. This increased activity during the pellet retrieval is reasonable when considering that this task component is more complex and involves increased use of the distal forelimb muscles. Therefore, the increased task-related neural activity during the pellet retrieval may represent specific or increased activation of neurons during distal movements as has been observed in premotor areas of non-human primates during tasks requiring distal forelimb movements [28, 29]. Additionally, while the lever press is indirectly associated with the food reward, the pellet retrieval is directly associated with the reward. As RFA has been shown to encode an expectation of reward [26], some of the observed increases in activity may be associated with the direct expectation of the food reward.

The combination of gross and fine reaching movements into a single behavioral task provides a new tool to examine the neural basis of reaching and grasping in rodents. Previous studies have argued for a specific role for RFA in grasping based upon the patterns of movements elicited by long-train ICMS [30]. However, lesion studies have produced conflicting results with some studies showing deficits in reach-to-grasp tasks after a lesion to CFA [10, 17] while other studies have found grasp-specific deficits isolated to lesions of RFA and not CFA when temporary lesions were induced with cortical cooling [30]. Isolating the specific differences in the roles that RFA and CFA play in controlling reaching and grasping is complicated by the fact that both regions exhibit extensive task-related neural activity during reach-to-grasp tasks [7] that are likely affected in part by the complex interactions between the two regions [15]. Our data have found that task-related activity in RFA was most modulated when aligned to the onset of reaching movements, however this may be due to a role of RFA in planning the combined reach-to-grasp movement as opposed to a specific role of RFA in reaching as opposed to grasping.

Because rodents have both primary and secondary forelimb motor regions, a number of studies have utilized rodent models of stroke and traumatic brain injury to examine the role of neuroplasticity in motor recovery [9, 10, 17]. In particular, ICMS mapping studies have found that expansions of RFA are associated with recovery of function following a lesion to CFA [10]. While rodents can learn to modulate activity in the perilesional cortex [6], the specific relationship between functional changes in perilesional cortex and secondary motor regions is unknown. The combined behavioral task demonstrated here is particularly relevant for examining the changes in task-related neural activity associated with post-injury neuroplasticity. While the lever press requires only a gross reaching movement, the pellet retrieval has the additional requirements of fine control of the distal forepaw and increased integration of sensory and motor information to successfully grasp and retrieve the pellet. Furthermore, the lever press task is more likely to be successfully completed in the early period of motor recovery which will allow for the assessment of the contribution of neuroplasticity to changes in task-related neural responses during the subacute period when the level of motor recovery is insufficient to allow for successful performance on the single-pellet retrieval task [9, 18].

Importantly, the task-related changes in neural activity were stable in individual channels over the 7-10 week period examined. While both task-related multi-unit firing and high-gamma band power changes persist over this period, the LFP high-gamma band showed more consistency across days. This increased stability is likely due to the difficulty in isolating the same individual neurons across days and the increased susceptibility of multi-unit firing rates to noise in the recordings. Previous studies examining chronic recordings in humans and non-human primates have observed that less than 40% of single-units were stable through a period of 15 days [31]. Additionally, changes in firing rate and spike amplitude are even seen within single recording days [32]. The finding that reaching related neural activity in RFA and CFA is stable over several weeks has particularly important implications for examining the role of neuroplasticity in recovery from a focal cortical lesion. Specifically, while multi-unit firing may be used to compare the proportion of channels with firing rate changes aligned to specific components of the task with greater temporal precision, LFP signals may be more valuable for examining changes in the relative strength of activations across days in single channels.

## 5. Conclusions

Collectively, we have demonstrated a novel complex rodent reaching task incorporating a gross lever press with a fine pellet retrieval into a single trial. Importantly, there were widespread task-related changes in neural activity during both task periods from microelectrodes in RFA and CFA, demonstrating a significant cortical involvement in both reaching tasks. Furthermore, this cortical involvement is maintained over months with stable task-related changes in neural activity observed at the level of single channels. These results serve to further characterize the normal role of rodent primary and secondary motor areas in planning and executing forelimb movements and further establish the rat as a model species for future studies examining the changes in the specific relationship between neural activity and forelimb movements associated with neuroplasticity following manipulation to the rodent sensorimotor system.

## Author Contributions

Conceived and designed the experiments: DTB DJG RJN; Performed the experiments: DTB DJG MDM; Analyzed the data: DTB; Contributed analysis tools: MDM; Wrote the paper: DTB RJN

## Acknowledgments

We would like to thank Mairaj Sami and Daniel Rittle for assistance with rodent behavioral training. Funding for this work was provided by the National Institutes of Health: NIH Grant R01NS030853, NIH Grant F32NS100339, and NIH Grant T32HD057850.

## Competing Interests

The authors have no relevant financial conflicts of interest related to this work.

## Data Availability

The data underlying the findings reported in this manuscript have been deposited in an online Open Science Framework repository that will be made public at the time of publication in accordance with the journal Data Availability Policy.

## References

1. Igarashi J, Isomura Y, Arai K, Harukuni R, Fukai T. A theta-gamma oscillation code for neuronal coordination during motor behavior. J Neurosci. 2013;33(47):18515–30. doi: 10.1523/JNEUROSCI.2126-13.2013. PubMed PMID: 24259574.

2. Slutzky MW, Jordan LR, Bauman MJ, Miller LE. A new rodent behavioral paradigm for studying forelimb movement. J Neurosci Methods. 2010;192(2):228–32. Epub 2010/08/10. doi: 10.1016/j.jneumeth.2010.07.040. PubMed PMID: 20691727; PubMed Central PMCID: PMCPMC2943042.

3. Hermer-Vazquez L, Hermer-Vazquez R, Chapin JK. The reach-to-grasp-food task for rats: a rare case of modularity in animal behavior? Behav Brain Res. 2007;177(2):322–8. doi: 10.1016/j.bbr.2006.11.029. PubMed PMID: 17207541; PubMed Central PMCID: PMCPMC1885543.

4. Whishaw IQ, Pellis SM. The structure of skilled forelimb reaching in the rat: a proximally driven movement with a single distal rotatory component. Behav Brain Res. 1990;41(1):49–59. Epub 1990/12/07. PubMed PMID: 2073355.

5. Chapin JK, Moxon KA, Markowitz RS, Nicolelis MA. Real-time control of a robot arm using simultaneously recorded neurons in the motor cortex. Nat Neurosci. 1999;2(7):664–70. doi: 10.1038/10223. PubMed PMID: 10404201.

6. Gulati T, Won SJ, Ramanathan DS, Wong CC, Bodepudi A, Swanson RA, et al. Robust neuroprosthetic control from the stroke perilesional cortex. J Neurosci. 2015;35(22):8653–61. doi: 10.1523/JNEUROSCI.5007-14.2015. PubMed PMID: 26041930.

7. Hyland B. Neural activity related to reaching and grasping in rostral and caudal regions of rat motor cortex. Behav Brain Res. 1998;94(2):255–69. PubMed PMID: 9722277.

8. Laubach M, Wessberg J, Nicolelis MA. Cortical ensemble activity increasingly predicts behaviour outcomes during learning of a motor task. Nature. 2000;405(6786):567–71. doi: 10.1038/35014604. PubMed PMID: 10850715.

9. Nishibe M, Barbay S, Guggenmos D, Nudo RJ. Reorganization of motor cortex after controlled cortical impact in rats and implications for functional recovery. J Neurotrauma. 2010;27(12):2221–32. doi: 10.1089/neu.2010.1456. PubMed PMID: 20873958; PubMed Central PMCID: PMCPMC2996815.

10. Nishibe M, Urban ET, 3rd, Barbay S, Nudo RJ. Rehabilitative training promotes rapid motor recovery but delayed motor map reorganization in a rat cortical ischemic infarct model. Neurorehabil Neural Repair. 2015;29(5):472–82. doi: 10.1177/1545968314543499. PubMed PMID: 25055836; PubMed Central PMCID: PMCPMC4303553.

11. Slutzky MW, Jordan LR, Lindberg EW, Lindsay KE, Miller LE. Decoding the rat forelimb movement direction from epidural and intracortical field potentials. J Neural Eng. 2011;8(3):036013. doi: 10.1088/1741-2560/8/3/036013. PubMed PMID: 21508491; PubMed Central PMCID: PMCPMC3124348.

12. Neafsey EJ, Sievert C. A second forelimb motor area exists in rat frontal cortex. Brain Res. 1982;232(1):151–6. Epub 1982/01/28. PubMed PMID: 7055691.

13. Rouiller EM, Moret V, Liang F. Comparison of the connectional properties of the two forelimb areas of the rat sensorimotor cortex: support for the presence of a premotor or supplementary motor cortical area. Somatosens Mot Res. 1993;10(3):269–89. Epub 1993/01/01. PubMed PMID: 8237215.

14. Sievert CF, Neafsey EJ. A chronic unit study of the sensory properties of neurons in the forelimb areas of rat sensorimotor cortex. Brain Res. 1986;381(1):15–23. Epub 1986/08/27. PubMed PMID: 3530375.

15. Deffeyes JE, Touvykine B, Quessy S, Dancause N. Interactions between rostral and caudal cortical motor areas in the rat. J Neurophysiol. 2015;113(10):3893–904. doi: 10.1152/jn.00760.2014. PubMed PMID: 25855697; PubMed Central PMCID: PMCPMC4480625.

16. Whishaw IQ, Alaverdashvili M, Kolb B. The problem of relating plasticity and skilled reaching after motor cortex stroke in the rat. Behav Brain Res. 2008;192(1):124–36. doi: 10.1016/j.bbr.2007.12.026. PubMed PMID: 18282620.

17. Whishaw IQ, Pellis SM, Gorny BP, Pellis VC. The impairments in reaching and the movements of compensation in rats with motor cortex lesions: an endpoint, videorecording, and movement notation analysis. Behav Brain Res. 1991;42(1):77–91. PubMed PMID: 2029348.

18. Guggenmos DJ, Azin M, Barbay S, Mahnken JD, Dunham C, Mohseni P, et al. Restoration of function after brain damage using a neural prosthesis. Proc Natl Acad Sci U S A. 2013;110(52):21177–82. doi: 10.1073/pnas.1316885110. PubMed PMID: 24324155; PubMed Central PMCID: PMCPMC3876197.

19. Quiroga RQ, Nadasdy Z, Ben-Shaul Y. Unsupervised spike detection and sorting with wavelets and superparamagnetic clustering. Neural Comput. 2004;16(8):1661–87. Epub 2004/07/02. doi: 10.1162/089976604774201631. PubMed PMID: 15228749.

20. Marple Jr SL, Carey WM. Digital spectral analysis with applications. ASA; 1989.

21. Rasch MJ, Gretton A, Murayama Y, Maass W, Logothetis NK. Inferring spike trains from local field potentials. J Neurophysiol. 2008;99(3):1461–76. doi: 10.1152/jn.00919.2007. PubMed PMID: 18160425.

22. Ellens DJ, Gaidica M, Toader A, Peng S, Shue S, John T, et al. An automated rat single pellet reaching system with high-speed video capture. J Neurosci Methods. 2016;271:119–27. doi: 10.1016/j.jneumeth.2016.07.009. PubMed PMID: 27450925; PubMed Central PMCID: PMCPMC5003677.

23. Wong CC, Ramanathan DS, Gulati T, Won SJ, Ganguly K. An automated behavioral box to assess forelimb function in rats. J Neurosci Methods. 2015;246:30–7. Epub 2015/03/15. doi: 10.1016/j.jneumeth.2015.03.008. PubMed PMID: 25769277; PubMed Central PMCID: PMCPMC5472046.

24. Jensen W, Rousche PJ. Encoding of self-paced, repetitive forelimb movements in rat primary motor cortex. Conf Proc IEEE Eng Med Biol Soc. 2004;6:4233–6. doi: 10.1109/IEMBS.2004.1404180. PubMed PMID: 17271238.

25. Khorasani A, Heydari Beni N, Shalchyan V, Daliri MR. Continuous Force Decoding from Local Field Potentials of the Primary Motor Cortex in Freely Moving Rats. Sci Rep. 2016;6:35238. Epub 2016/10/22. doi: 10.1038/srep35238. PubMed PMID: 27767063; PubMed Central PMCID: PMCPMC5073334.

26. Saiki A, Kimura R, Samura T, Fujiwara-Tsukamoto Y, Sakai Y, Isomura Y. Different modulation of common motor information in rat primary and secondary motor cortices. PLoS One. 2014;9(6):e98662. doi: 10.1371/journal.pone.0098662. PubMed PMID: 24893154; PubMed Central PMCID: PMCPMC4043846.

27. Poddar R, Kawai R, Olveczky BP. A fully automated high-throughput training system for rodents. PLoS One. 2013;8(12):e83171. doi: 10.1371/journal.pone.0083171. PubMed PMID: 24349451; PubMed Central PMCID: PMCPMC3857823.

28. Kurata K, Tanji J. Premotor cortex neurons in macaques: activity before distal and proximal forelimb movements. J Neurosci. 1986;6(2):403–11. Epub 1986/02/01. PubMed PMID: 3950703.

29. Rizzolatti G, Camarda R, Fogassi L, Gentilucci M, Luppino G, Matelli M. Functional organization of inferior area 6 in the macaque monkey. II. Area F5 and the control of distal movements. Exp Brain Res. 1988;71(3):491–507. Epub 1988/01/01. PubMed PMID: 3416965.

30. Brown AR, Teskey GC. Motor cortex is functionally organized as a set of spatially distinct representations for complex movements. J Neurosci. 2014;34(41):13574–85. doi: 10.1523/JNEUROSCI.2500-14.2014. PubMed PMID: 25297087.

31. Dickey AS, Suminski A, Amit Y, Hatsopoulos NG. Single-unit stability using chronically implanted multielectrode arrays. J Neurophysiol. 2009;102(2):1331–9. doi: 10.1152/jn.90920.2008. PubMed PMID: 19535480; PubMed Central PMCID: PMCPMC2724357.

32. Perge JA, Homer ML, Malik WQ, Cash S, Eskandar E, Friehs G, et al. Intra-day signal instabilities affect decoding performance in an intracortical neural interface system. J Neural Eng. 2013;10(3):036004. doi: 10.1088/1741-2560/10/3/036004. PubMed PMID: 23574741; PubMed Central PMCID: PMCPMC3693851.

